# Global analysis of suppressor mutations that rescue human genetic defects

**DOI:** 10.1101/2022.11.09.515781

**Authors:** Betül Ünlü, Carles Pons, Uyen Linh Ho, Amandine Batté, Patrick Aloy, Jolanda van Leeuwen

**Author notes:** These authors contributed equally to this work (Betül Ünlü, Carles Pons).

## Abstract

Genetic suppression occurs when the deleterious effects of a primary “query” mutation, such as a disease-causing mutation, are rescued by a suppressor mutation elsewhere in the genome. To capture existing knowledge on suppression relationships between human genes, we examined 2,400 published papers for potential interactions identified through either genetic modification of cultured human cells or through association studies in patients. The resulting network encompassed 476 unique suppression interactions that frequently linked genes that function in the same biological process. Suppressors were strongly enriched for genes with a role in stress response or signaling, suggesting that deleterious mutations can often be buffered by modulating signaling cascades or immune responses. Suppressor mutations tended to be deleterious when they occurred in absence of the query mutation, in apparent contrast with their protective role in the presence of the query. We formulated and quantified mechanisms of genetic suppression that could explain 71% of interactions and provided mechanistic insight into disease pathology. Finally, we used these observations to accurately predict suppressor genes in the human genome. The emerging frequency of suppression interactions and range of underlying mechanisms suggest that compensatory mutations may exist for the majority of genetic diseases.

## BACKGROUND

Despite our progress in sequencing genomes, translating the variants detected in an individual into knowledge about disease risk or severity remains challenging. The relationship between genotype and phenotype is complex because genes and their products function as components of dynamic networks, with each gene or protein linked to many others through genetic and physical interactions. Modifying mutations in such interaction partners can either increase the severity of a genetic trait, or can have a protective effect and compensate for the deleterious effects of a particular mutation, a phenomenon referred to as genetic suppression [1, 2]. Genetic suppression is of particular interest for human disease, as suppressors of disease alleles highlight biological mechanisms of compensation, thereby potentially uncovering new therapeutic strategies. For example, a genome-wide association study discovered a loss-of-function variant in *BCL11A*, encoding a transcriptional repressor of fetal hemoglobin subunit γ, as protective against severe β-thalassemia [3]. When expressed in adults, the γ -subunit of hemoglobin can replace the β-subunit, which is mutated in β-thalassemia patients, a finding that led to the development of gene therapies targeting *BCL11A* [4]. Despite its success, this approach for discovering protective modifiers cannot be universally applied, as most monogenic diseases and/or protective variants are too rare for such systematic association studies [1]. Alternative methods to identify suppressor genes are thus needed.

The systematic mapping of large numbers of suppressor mutations can highlight properties of suppression interactions that can be used to find or predict suppressors in other contexts [5]. To date, such systematic analyses have only been performed in inbred model organisms, enabling the rigorous assessment of the effects of combining mutations in an otherwise isogenic background. These systematic suppression studies have led to the discovery of specific mechanistic classes of suppression [6-9]. In bacteria, fungi, fly, and worm, most extragenic suppression interactions occur between genes that are annotated to the same biological process [8, 10-14]. Extragenic suppression of partial loss-of-function alleles can also occur through general mechanisms of suppression, which are often allele-specific and affect the translation of the deleterious mutation, the expression of the affected gene, or the stability of its gene-product [6, 7, 15]. Together, these mechanisms of suppression explain ∼70% of all described suppression interactions in the budding yeast *Saccharomyces cerevisiae* [15] and have been used to predict suppressors among hundreds of genes on aneuploid chromosomes [5].

Aside from the mechanisms of suppression that have been described in model organisms, additional biological mechanisms of compensation may exist in humans that could be of relevance for understanding variation in disease severity or penetrance. Here, we systematically analyzed suppression interactions among human genes to define general principles of suppression specific to humans. A thorough understanding of suppression mechanisms and properties may guide the discovery or prediction of protective alleles for rare genetic diseases, which could direct the rational design of new therapeutics.

## RESULTS

### A network of literature-curated suppression interactions

To identify and annotate existing suppression interactions among human genes, we examined 2,400 published papers for potential interactions **(Data S1**). Papers were derived from the BioGRID [16], OMIM [17], specific PubMed searches (see Methods), and references found within the examined papers. We considered suppression interactions from two types of studies. First, we included interactions identified through genetic modifications in cultured human cells. Two genes were considered to have a suppression interaction when the genetic perturbation of a “query” gene led to reduced survival, decreased proliferation, or was otherwise associated with decreased cellular health, which was rescued by mutation of a different gene (the “suppressor” gene).

Second, we included interactions found through association studies in patients. Two genes were considered to have a suppression interaction when the disease risk or severity associated with a particular allele of a query gene was reduced in the presence of a minor allele of a suppressor gene. We excluded cancer patients in our analysis, as cancer is a disease of increased cell proliferation and thus mechanistically quite different from diseases caused by decreased cellular health. For genome-wide association studies (GWAS), we generally considered the gene that was closest to the SNP with the most significant association to a protective effect to be the suppressor gene. While the gene that is closest to a GWAS peak is not always the causal gene, it is in about 70-80% of the cases [18-21].

When data was provided supporting that another gene was driving the suppression phenotype, we based our suppressor annotation on this additional evidence (see Methods). For both cell-derived and patient-derived interactions, we excluded suppression interactions that were intragenic (occurring between two mutations within the same gene), occurred between more than two genes, or involved the major allele of either the query or the suppressor gene from the final dataset.

In total, we collected 932 suppression interactions from 466 papers. From each interaction, we annotated the system in which the interaction was identified (cultured cells or patients), the query and suppressor mutations and whether these had a loss- or gain-of-function effect, the used cell line or affected tissue, the relative effect size of the suppression, whether any drugs were used, and the disease (if applicable). After removing duplicate interactions that had been described multiple times, the resulting network encompassed 476 unique suppression interactions for 93 different query genes (**Fig. 1A**). Four interactions were identified in both directions, such that both suppressor and query mutations were deleterious, but combination of the two gene mutants could restore fitness (**Data S2**). In total, 302 unique interactions were identified in cultured cells and 180 in patients (**Fig. 1B**). Although we observed significant overlap between these two subnetworks (6 shared interactions, p<0.0005, Fisher’s exact test), 99% of interactions were reported in only one type of study (either in cultured cells or in patients).

**Figure 1.**
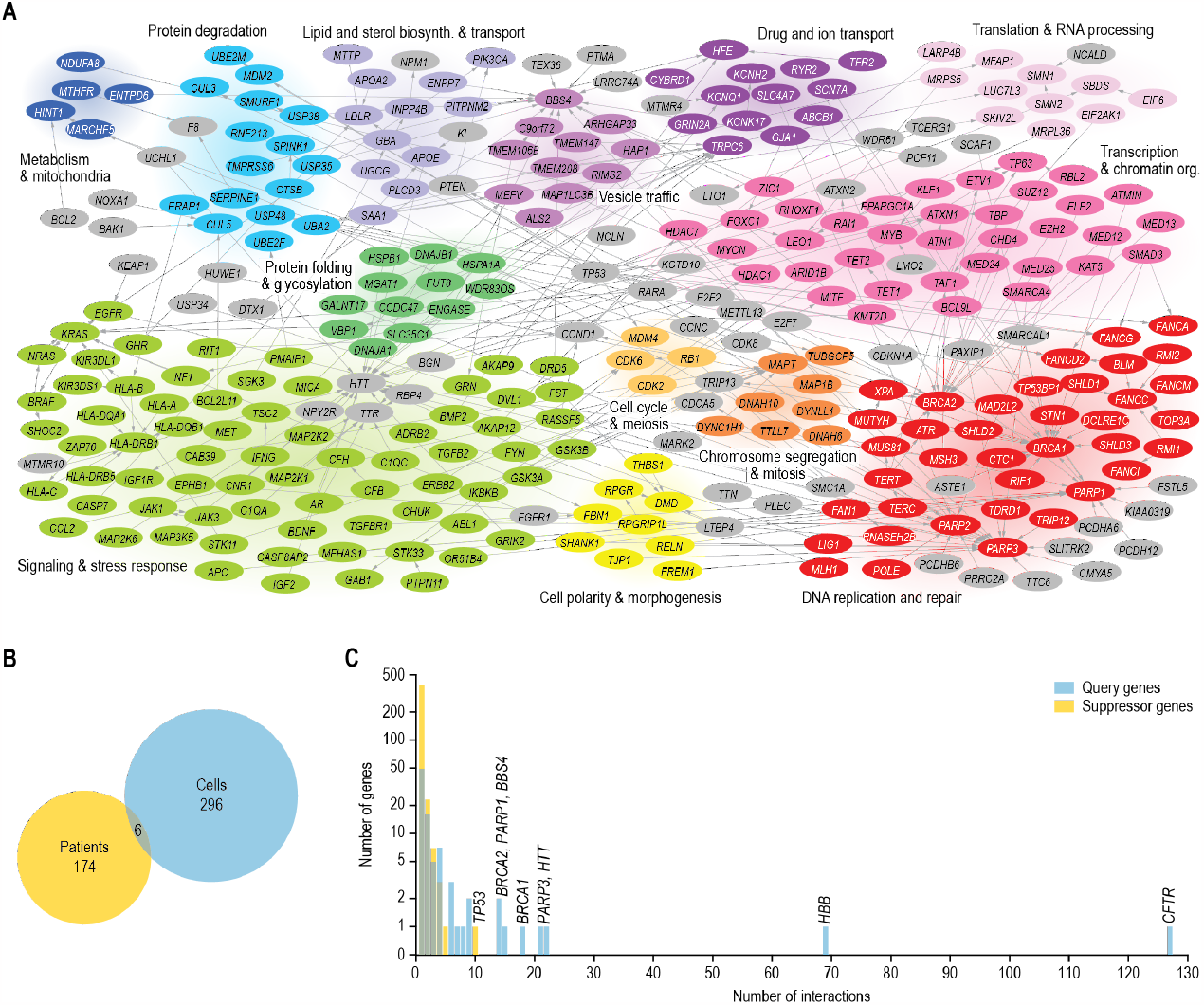
A literature-curated suppression network. (**A**) A network of suppression interactions curated from the biomedical literature. Suppression interactions are represented as arrows that point from the suppressor to the query gene. Nodes are colored and grouped based on the function of the gene. Gray nodes indicate genes that are poorly characterized, multifunctional, or have functions that are not otherwise represented in the figure. For clarity, interactions involving *HBB* and *CFTR*, which have a high interaction degree, are not shown in the figure. (**B**) Proportional Venn diagram showing the number of unique suppression interactions that have been identified in cultured cells, in patients, or in both. (**C**) Degree distribution of suppression interactions. The number of unique suppression interactions is plotted against the number of query or suppressor genes showing that number of interactions.

The vast majority of suppressor genes (92%) suppressed a single query gene (**Fig. 1C, Data S2**). The most common suppressor gene, *TP53*, interacted with 10 queries. The encoded protein, p53, induces cell cycle arrest and apoptosis in response to various stresses [22] and the suppressed query genes are functionally diverse with roles in transcription (*TP63*), DNA repair (*FANCA, FANCD2, FANCG*), protein degradation (*CUL3, UBE2M, KCTD10*), ribosome maturation (*SBDS*), and p53 regulation (*MDM2, MDM4*). Although loss of p53 can cause uncontrolled cell proliferation and tumor formation, heterozygous mutation of *TP53* can be beneficial under conditions that would otherwise lead to excessive cell death. For example, mutation of a single copy of *TP53* can protect against severe bone marrow failure in patients with Shwachman-Diamond syndrome [23]. In contrast to the low interaction degree observed for most suppressors, about half of the query genes (46%) were suppressed by multiple suppressor genes, with eight query genes (*BBS4, BRCA1, BRCA2, CFTR, HBB, HTT, PARP1*, and *PARP3*) interacting with more than 10 suppressors (**Fig. 1C**). Especially for *CFTR* (127) and *HBB* (69), high numbers of suppressor genes have been described, likely because mutations in these genes lead to relatively common Mendelian disorders resulting in the availability of rather large numbers of patients to study. We excluded interactions of *CFTR* and *HBB* from the analyses described in the following sections, to prevent potential bias of our results by the high number of interactions described for these genes.

### Suppressor genes are essential for optimal health and cellular fitness

Consistent with their requirement for maintaining health or cellular fitness, query genes were significantly more likely to be intolerant to loss-of-function mutation in the human population, had a more deleterious effect on the proliferation of cultured human cells when inactivated, and tended to be conserved in a higher number of species than other genes in the human genome (**Fig. 2A-C**). In general, query genes that were suppressed in cellular models had more severe phenotypes than those described in patients (**S1A-C**). In apparent contrast with their role in ameliorating phenotypes in the presence of the query mutation, suppressor genes were also significantly depleted for deleterious mutations in the human population, were generally required for optimal proliferation of cultured cells, and tended to be highly conserved across species (**Fig. 2A-C, S1A-C**). Furthermore, mutations in suppressor genes were often associated with diseases themselves (**Fig. 2D, S1D**). The deleteriousness of query and suppressor mutations was weakly correlated (**Fig. S1E**). These results suggest that the beneficial effects of suppressor mutations may only be apparent in the presence of the query mutation. Alternatively, because these analyses look at the effect of deleterious mutations in the suppressor gene, the variants that cause the suppression phenotype may not lead to loss-of-function of the suppressor. To investigate the latter possibility, we considered gain-of-function and loss-of-function suppressor mutations separately **(Fig. 2E, S1F**). We did not observe significant differences in loss-of-function intolerance between genes carrying gain-of-function or loss-of-function suppressor mutations (**Fig. S1G**). Furthermore, when focusing solely on suppressor genes that were identified using knockout experiments in cell culture, 86% of these genes were needed for optimal cellular proliferation. These results show that the loss-of-function intolerance of suppressor genes cannot be explained by a preference for gain-of-function suppressor mutations in these genes. Suppressor mutations thus appear to be frequently detrimental in the absence of the query mutation.

**Figure 2.**
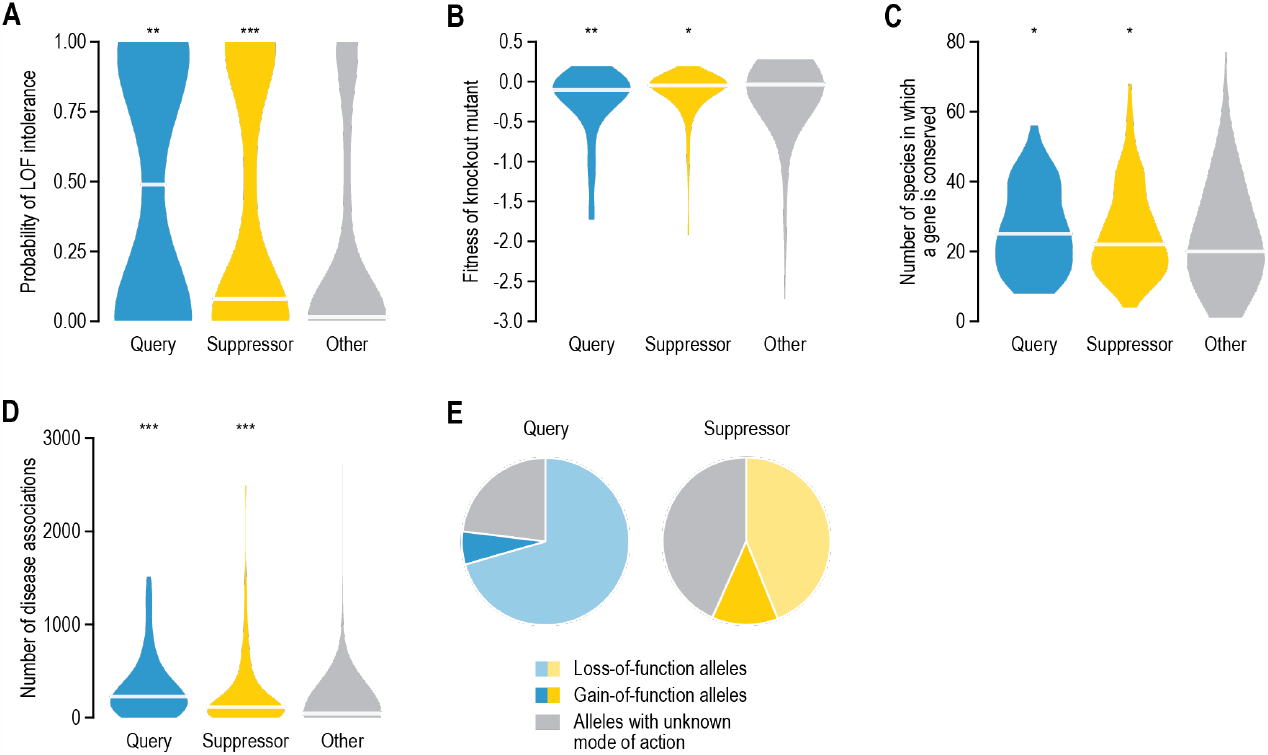
Suppressor genes are important for maintaining health and cellular fitness. (**A**) Probability of loss-of-function intolerance for query genes, suppressor genes, and all other genes, based on the frequency of deleterious variants affecting the genes in the human population [24]. (**B**) Median effect of gene knockout on cell proliferation determined as the change in abundance of guide RNAs targeting a gene in pooled CRISPR-Cas9 screens across 1,070 cell lines [25], for the same gene groups as in (A). (**C**) Number of species in which an ortholog of the query or suppressor gene is present. (**D**) The number of diseases that are associated with a gene in DisGeNET [26], for the same gene groups as in (A). (**E**) The fraction of query and suppressor genes that have loss-of-function, gain-of-function, or unknown modes of action. Statistical significance compared to the “Other” group was determined using Mann-Whitney U tests. * p<0.05, ** p<0.005, *** p<0.0005. Horizontal lines in violin plots: median.

### Overlap with other interaction networks

The suppression interactions overlapped significantly with protein-protein interactions and various types of genetic interactions (**Fig. S2**) [16]. Positive genetic interactions occur when a defect of a double mutant is less severe than expected based on the phenotypes of the single mutants [27]. In contrast, negative genetic interactions, such as synthetic lethality, occur when the phenotype of a double mutant is more severe than expected [27]. The overlap between suppression interactions and positive genetic interactions is thus not surprising, as suppression interactions are an extreme type of positive interaction (**Fig. S2**). The overlap with negative genetic interactions reflects that mutations in a gene may lead to either loss-of-function or gain-of-function effects, which may display opposite types of genetic interactions (**Fig. S2**) [8]. Despite the overlap with other interaction networks, the vast majority of suppression interactions (80%) are specific to the suppression network and thus highlight novel functional connections between genes.

### Suppression interactions within and across cellular processes

Consistent with other organisms [8, 13-15], suppression interactions in human often occurred between functionally related genes, such that a query mutant tended to be suppressed by another gene annotated to the same biological process (**Fig. 3A, S3A**). Genes connected by suppression interactions also tended to be co-expressed and encode proteins that function in the same subcellular compartment and/or belong to the same pathway or protein complex (**Fig. 3B**). The extent of functional relatedness between suppression gene pairs did not depend on the conditions under which the interaction was identified (e.g., in the presence of a specific drug), whether the interaction was discovered in patients or in cultured cells, the number of times a particular interaction had been described, the relative effect size of the suppression, or whether the mutations had a gain- or loss-of-function effect (**Fig. S3B**). When multiple suppressors had been described for a query gene across independent studies, the suppressor genes also tended to be co-expressed and encode proteins that function in the same pathway or protein complex and/or that localize to the same subcellular compartment, suggesting that the suppressor genes functioned through similar molecular mechanisms (**Fig. 3C**).

**Figure 3.**
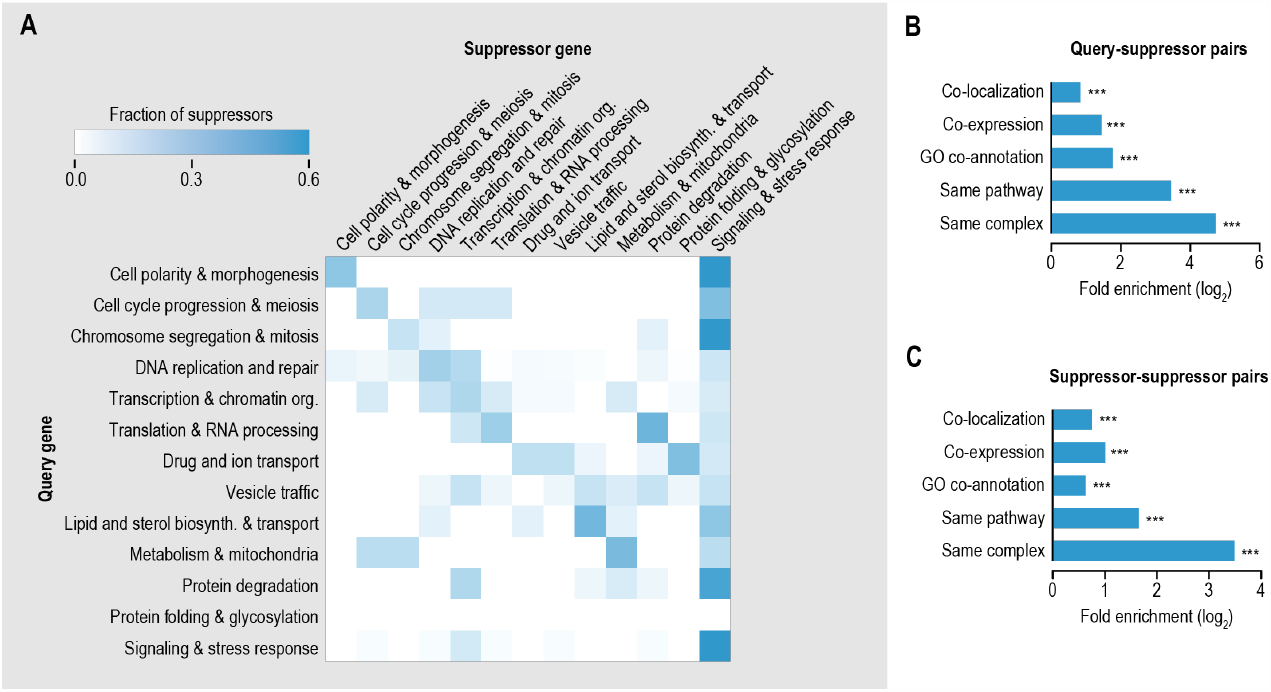
Functional connections between query and suppressor genes. (**A**) Frequency of suppression interactions connecting genes within and across indicated biological processes. Color intensity reflects the fraction of suppressor genes belonging to a particular biological process for all interactions involving query genes annotated to a given biological process. (**B**-**C**) Fold enrichment for co-localization, co-expression, GO co-annotation, same pathway membership, and same complex membership for query-suppressor gene pairs (B) or among suppressor genes that have been described for the same query gene (C).

Despite their tendency to connect functionally related genes, suppression interactions also linked different biological processes. Genes with a role in signaling or the response to stress suppressed defects associated with mutation of genes involved in many different biological processes. This central role for signaling and stress response in the suppression network was observed both for interactions identified in patients and for those found in cultured cells (**Fig. S3C**). The suppressor genes in this category often played a role in protein phosphorylation and kinase cascades (60%) and/or in apoptosis or its regulation (48%). Moreover, in patients with inflammatory diseases, such as multiple sclerosis, the suppressor genes frequently encoded members of the major histocompatibility complex family that play a critical role in the immune system [28].

Genes involved in chromatin organization or transcription were also strongly overrepresented as suppressors, mainly in interactions identified in cultured cells (**Fig. 3A, S3C**). These interactions reflect a mechanism whereby modified expression of genes encoding members of the same pathway as the query gene can compensate for the altered activity of the query. For example, the deleterious effect of loss of *BRCA2*, which encodes a protein with a role in double-strand DNA break repair via homologous recombination, can be rescued by silencing transcriptional repressor E2F7 [29]. E2F7 inhibits expression of several genes with a role in recombination or double-strand break repair, including *CHEK1, DMC1, GEN1*, and *MND1*, that when expressed can potentially compensate for the absence of BRCA2. In total, we found that ∼44% of suppressor genes that encode characterized transcription factors affect expression of query pathway members (see Methods).

### Mechanistic categories of suppression interactions

We classified the suppression interactions into distinct mechanistic categories on the basis of the functional relationship between the query and suppressor genes. In many of the reported interactions (33%), the query genes were suppressed by mutations in functionally related genes (“Functional mechanisms”, **Fig. 4A,C**). These include interactions in which both the query and the suppressor genes encode members of the same protein complex (“Same complex”, 6% of interactions) or pathway (“Same pathway”, 13% of interactions). Seven percent of interactions involved suppression by a different, but related, pathway or complex (“Alternative pathway”). In this scenario, the deleterious phenotype caused by absence of a specific function required for normal (cellular) health is suppressed when an alternative pathway is rewired to re-create the missing activity. Finally, 7% of gene pairs were annotated to the same biological process but pathway or complex annotation data were not available for one or both genes (“Uncharacterized functional connection”). In addition to suppression interactions between functionally related genes, more general, pleiotropic classes of suppressors exist that affect degradation of the mutated query protein or mRNA, gene expression, or signaling and stress response pathways (“General mechanisms”, **Fig. 4A,D**). Together, these general mechanisms of suppression explain 38% of interactions, with half of these (19%) involving altered signaling or stress response processes. In total, 71% of interactions could be assigned to a mechanistic class.

**Figure 4.**
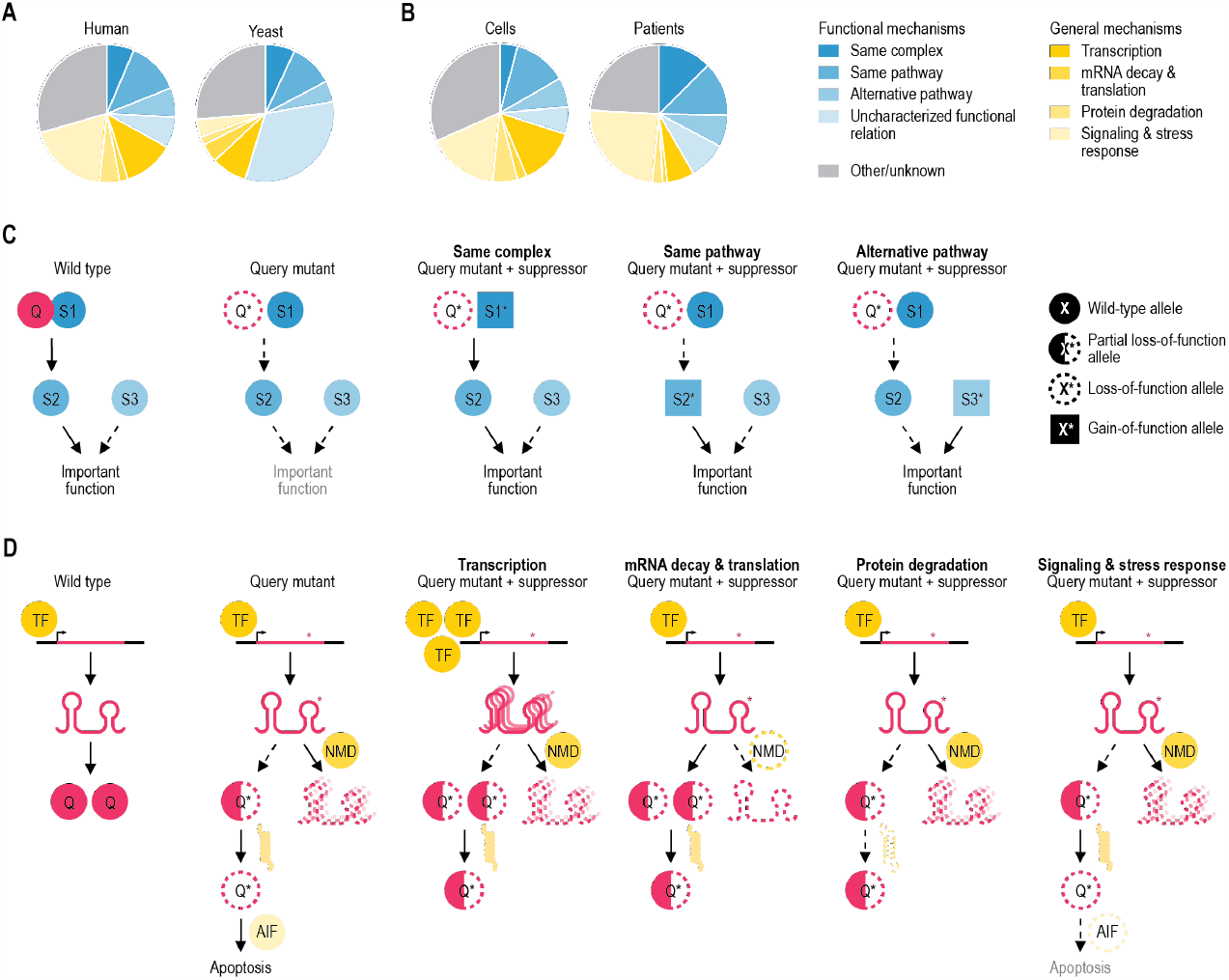
Mechanistic classes of suppression. (**A**) Distribution of suppression interactions across mechanistic classes for interactions identified in this study (left) or interactions described in the budding yeast using a similar literature curation approach (right) [8]. (**B**) Distribution of suppression interactions across mechanistic classes for interactions discovered in cultured human cells (left) or in patients (right). (**C**) Mechanisms of suppression between genes encoding proteins that function within the same biological process are illustrated. In a situation where the query (“Q”) activates a protein S2, which has an important biological function, suppression can take place in multiple ways. For example, the suppressor (S1) can be part of the same complex as the query, and gain-of-function mutations in S1 can restore the activation of S2. Alternatively, suppression can occur through gain-of-function mutations in S2, such that it no longer requires the query protein for its activation. The suppressor (S3) can also function in an alternative, but related, pathway. Specific alterations in this alternative pathway can restore the important function that was lacking in the absence of the query protein. (**D**) General mechanisms of suppression among pairs of genes that do not share a close functional relationship are illustrated. Often, general suppression is associated with partial loss-of-function query alleles that carry mutations that destabilize the protein or mRNA, leading to deleterious phenotypes due to reduced levels of the query protein. Partial loss-of-function query alleles can be suppressed by increasing protein expression, for instance via increased transcription of the query gene or through decreased degradation of the mutant mRNA via mutation of the nonsense-mediated mRNA decay (NMD) pathway. Partial loss-of-function mutations can also be suppressed by inactivation of a member of the protein degradation pathway, which may expand the pool of partially functional query protein. Finally, suppression may occur through inhibition of apoptosis. TF: transcription factor; AIF: apoptosis-inducing factor.

When comparing suppression interactions described among human genes to those identified using a similar literature curation approach in the budding yeast *Saccharomyces cerevisiae* [8], there were significant differences in the distribution of the interactions across mechanistic classes (**Fig. 4A**). Notably, whereas 55% of suppression interactions in yeast occurred between genes with a functional connection, only 33% of the human suppression gene pairs were functionally related (p<0.0005 comparing yeast to human, Fisher’s exact test). Although the yeast genome is more extensively functionally annotated, this is unlikely to be the cause of this difference, as nearly all genes considered here have a biological process annotation (**Data S2**) and the percentage of unclassified gene pairs is similar between the two datasets (26% for yeast, 29% for human, p=0.31, Fisher’s exact test). In contrast, the percentage of gene pairs involving a general suppression mechanism, in particular suppression by modifying the stress response or signaling pathways, was significantly lower for yeast gene pairs compared to human suppression interactions (19% for yeast, 38% for human, p<0.0005, Fisher’s exact test).

The observed differences between yeast and human could be due to differences in the methods used to identify suppression interactions. Yeast suppressor isolation experiments generally rely on genetically engineered query mutant alleles, such as gene deletion alleles or temperature sensitive point mutants, and defined laboratory environments, whereas interactions detected in patients occur between natural variants in an uncontrolled setting. Because interactions that were discovered in cultured human cells also often involved genome modification and controlled laboratory environments, we investigated the distribution across mechanistic classes separately for interactions identified in cultured cells and those found in patients. The distribution across mechanistic classes between the two sets of human suppression interactions was largely similar (**Fig. 4B**). Although interactions found in patients more often involved suppression by altering signaling or stress response pathways than those in cultured cells, the percentage of interactions involving suppression by signaling or stress response genes was still significantly higher in cultured human cells than in yeast (p<0.0005, Fisher’s exact test). Moreover, the fraction of interacting gene pairs with a functional connection was lower in cultured cells compared to patients, in contrast to the high percentage of functionally related pairs seen for yeast (**Fig. 4A,B**). Thus, experimental factors do not appear to be the main cause of the observed differences in frequency of suppression mechanisms between yeast and human.

### Suppressors provide mechanistic insight into disease pathology

Combining data from multiple suppression studies can reveal the general significance of particular protein classes in attenuating disease phenotypes. As mentioned above, a relatively high number of suppressor genes have been identified for *HBB* and *CFTR*, which are mutated in sickle cell disease/β-thalassemia and cystic fibrosis patients, respectively (**Fig. 1C**). To investigate the molecular mechanisms driving suppression of these two query genes, we examined the 69 *HBB* and 127 CFTR suppressors in more detail, using our mechanistic suppressor classification (**Fig. 4**). Our systematic analysis highlighted both similarities and differences in disease pathology between the two diseases (**Fig. 5**).

**Figure 5.**
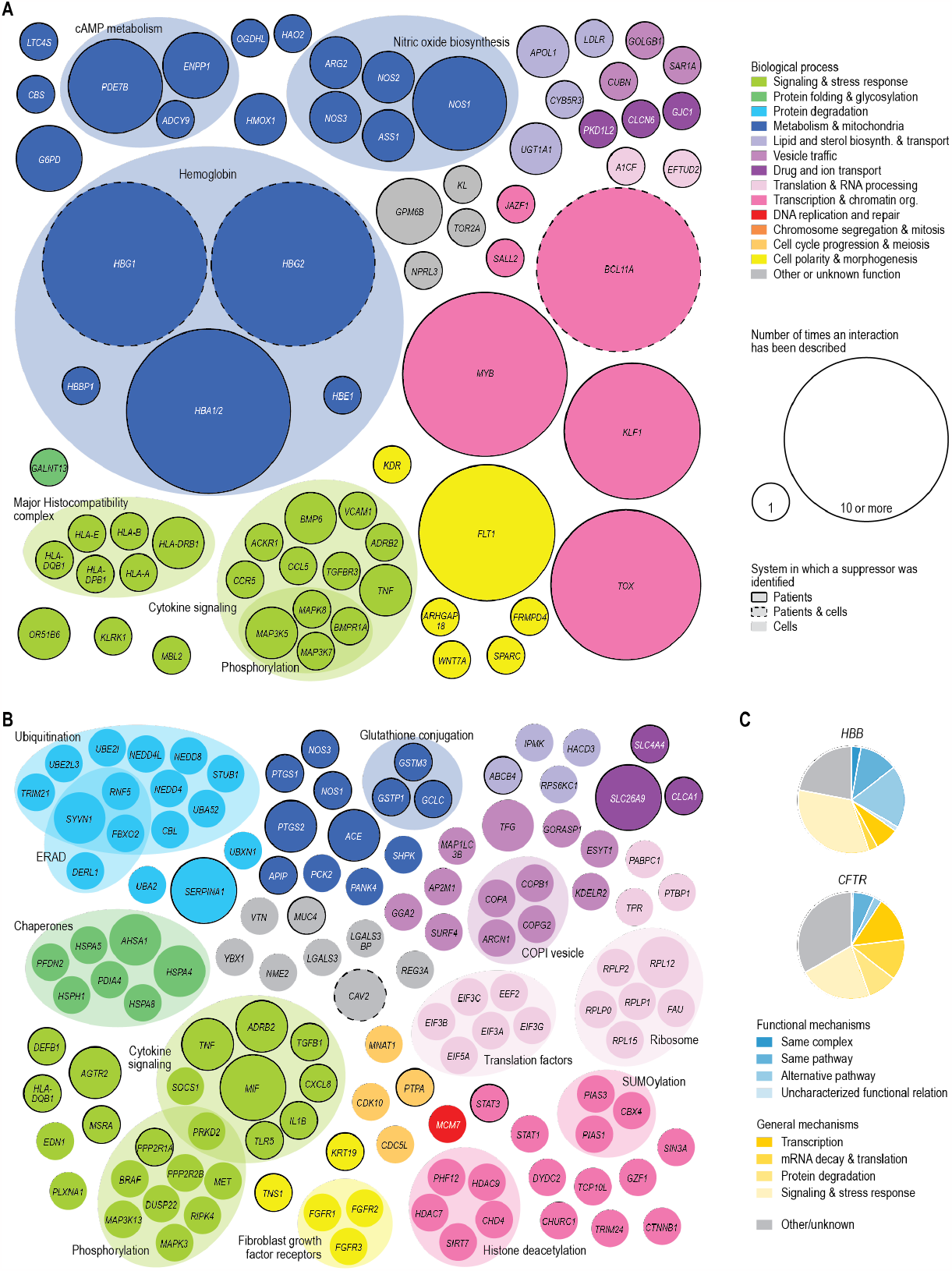
Suppressors of *HBB* and *CFTR* provide insight into disease pathology. (**A**-**B**) Suppressor genes that have been described for *HBB* (A) and *CFTR* (B). Nodes are colored and grouped based on the function of the genes. Gray nodes indicate genes that are poorly characterized, multifunctional, or that have functions that are not otherwise represented in the figure. Node size represents the number of times an interaction has been described. Nodes with a black border indicate suppressor genes that have been described in patients, nodes without a border represent suppressor genes that have been identified in cultured cells, and those with a dashed border have been found in both systems. (**C**) Distribution of *HBB* and *CFTR* suppression interactions across the mechanistic classes described in Fig. 4.

Attenuating cytokine signaling could for example reduce symptoms of both cystic fibrosis and sickle cell disease, highlighting the importance of inflammation in both cases (**Fig. 5**) [30, 31]. However, whereas *HBB* suppressors frequently occurred in genes with a functional connection to *HBB, CFTR* suppressors tended to function through more general mechanisms of suppression (**Fig. 5C**). The most commonly found suppressors of *HBB*, encoding the β-subunit of hemoglobin, encode either other hemoglobin subunits (i.e. *HBA1/2, HBG2*) or their transcriptional regulators (i.e. *BCL11A, MYB*) (**Fig. 5A,C**). These other hemoglobin subunits can either functionally replace the mutated β-subunit or balance the ratio of hemoglobin subunits, thereby increasing the relative amount of functional hemoglobin [32]. Thus, suppressors of complete loss-of-function mutations in *HBB* function through circumventing the need for HBB. In contrast, suppressors of *CFTR* mutants tend to restore CFTR function (**Fig. 5B,C**). *CFTR* encodes an ion channel located on the plasma membrane of epithelial cells where it regulates the flow of chloride and bicarbonate ions in and out of the cell. The F508del mutation, an inframe-deletion that removes the phenylalanine residue at position 508, occurs in ∼90% of cystic fibrosis patients [33]. Although CFTR-F508del retains substantial function, it is recognized by the ER quality control machinery as misfolded and is prematurely degraded [34]. Changes in CFTR transcription or translation, chaperone levels, activity of the protein degradation machinery, or efficiency of ER to plasma membrane trafficking can (partially) restore expression of the mutant CFTR protein at the plasma membrane and explain 53% of CFTR suppression interactions. These examples highlight how integrating data from tens to hundreds of papers can provide insight on the general mechanisms through which suppression of particular disease mutations can occur.

### Query-suppressor gene pairs are often co-mutated in tumor cells

Cancer cells generally have increased genome instability and reduced DNA repair, leading to the accumulation of hundreds to thousands of mutations, the majority of which are considered passenger mutations that do not favor tumor growth [35, 36]. Because loss-of-function mutations in query genes tend to have a negative effect on cell proliferation, we suspected that damaging passenger mutations affecting query genes would be more likely to persist in a tumor if they were accompanied by mutations in the corresponding suppressor gene(s). To test this hypothesis, we first examined gene fitness data from genome-scale CRISPR-Cas9 gene knockout screens across 1,070 cancer cell lines from the Cancer Dependency Map (DepMap) project [25]. We found that knockout of the query genes led to more variable effects on cell proliferation than knockout of other genes that had a comparable mean fitness defect (**Fig. S4A**,**B**). This suggests that the deleterious consequences of loss of the query gene are buffered in some cell lines but not in others, potentially due to differences in the presence of suppressor variants. To further explore this possibility, we looked at the presence of damaging mutations in query and suppressor genes across 1,758 cancer cell lines [37]. We found that damaging mutations in the query gene were more frequently accompanied by mutations in the corresponding suppressor genes than expected by chance (**Fig. 6A**). Furthermore, we examined the co-occurrence of mutations in tumor samples collected from 69,223 patients across 213 different studies [38]. Also in these patient samples, impactful mutations in query genes frequently co-occurred with mutations in the corresponding suppressor genes (**Fig. 6B**). These results suggest that the suppressor mutations that lead to improved health of patients with a genetic disease or increased proliferation of cultured cells also provide a selective advantage to tumor cells carrying mutations in the same query gene.

**Figure 6.**
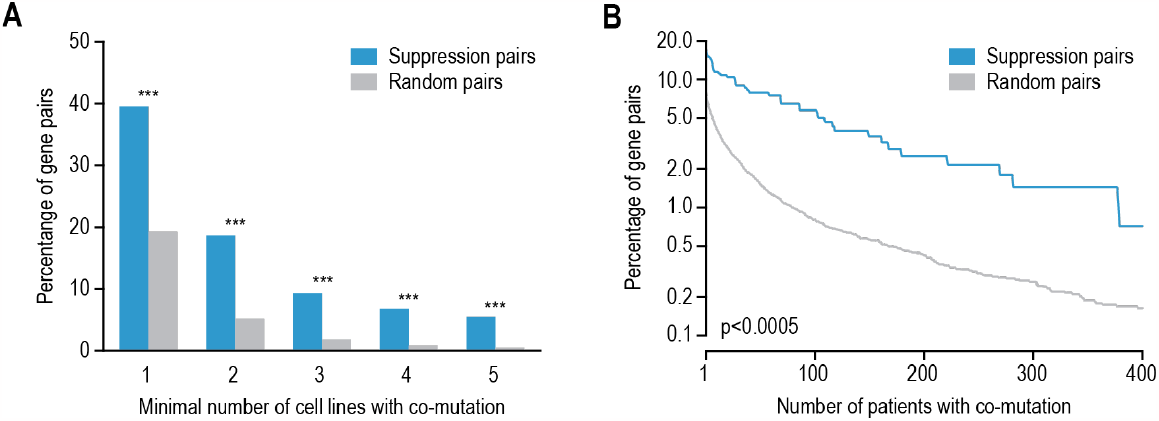
Suppression gene pairs are enriched for co-mutation in cancer samples. (**A**-**B**) Mutation co-occurrence in query and suppressor gene pairs in cancer cell lines (A) and in tumor samples obtained from patients (B) for gene pairs showing a suppression interaction or for randomized gene pairs. The percentage of gene pairs that were mutated in the same cell line or tumor sample is plotted against the number of cell lines or patient samples in which co-mutation was observed. Mutation data were obtained from the Cancer Dependency Map for cancer cell lines [37] and cBioPortal for patient samples [38]. Statistical significance was determined using Fisher’s exact test (A) or a Mann-Whitney U test (B). * p<0.05, ** p<0.005, *** p<0.0005.

### Predicting suppressor genes

Given the strong functional connection frequently observed between interacting query and suppressor genes (**Fig. 3**), we developed models that use these signatures to identify suppressor genes for a given query gene of interest (see Methods). First, we adapted a model we developed previously to predict suppressor genes in yeast [5] to the prediction of suppressors among human genes. In brief, this model scores and ranks potential suppressor genes by prioritizing close functional connections to the query gene. In this functional prioritization model, shared complex or pathway membership weigh more heavily than more distant functional connections, such as co-localization or co-expression. We used this suppressor prediction approach to identify candidate suppressor genes for all 93 query genes present in our dataset, by ranking all genes in the genome by their predicted likeliness of being a suppressor. For 25 query genes (27%), at least one validated suppressor gene ranked among the top 100 of those predicted, with 15 suppressor genes ranking in the top 10 (**Fig. S5, Data S3**). Consistent with the design of the model, 14 out of the 15 suppressors that were predicted with high accuracy encoded members of the same protein complex as the query gene.

Next, we aimed to further improve this model. We used a set of diverse features, including functional relationships (**Fig. 3**), other types of genetic and physical interactions (**Fig. S2**), and co-mutation in cancer cell lines (**Fig. 6**) to train a random forest classifier (see Methods). The random forest showed increased predictive power over the functional prioritization model, with 39 validated suppressor genes ranking among the top 100 of those predicted (**Fig. 7A, Data S4**). Only two suppressors would be expected to rank in the top 100 by random selection. In addition to predicting suppression interactions among genes with shared complex or pathway membership, the random forest model also accurately predicted 11 interactions involving genes with more distal functional relationships or general suppression mechanisms. For example, pathogenic variants of *MAPT*, encoding tau, can cause tau to aggregate, causing a range of neurodegenerative diseases. Suppression of *MAPT* by mutation of *GSK3A* or *GSK3B*, which encode kinases that hyperphosphorylate tau leading to its aggregation [39], was correctly predicted by the model. These results show that for at least 42% of query genes, the various properties that are generally observed for query-suppressor gene pairs can be used to narrow the search space for potential suppressor genes from thousands to less than a hundred genes.

**Figure 7.**
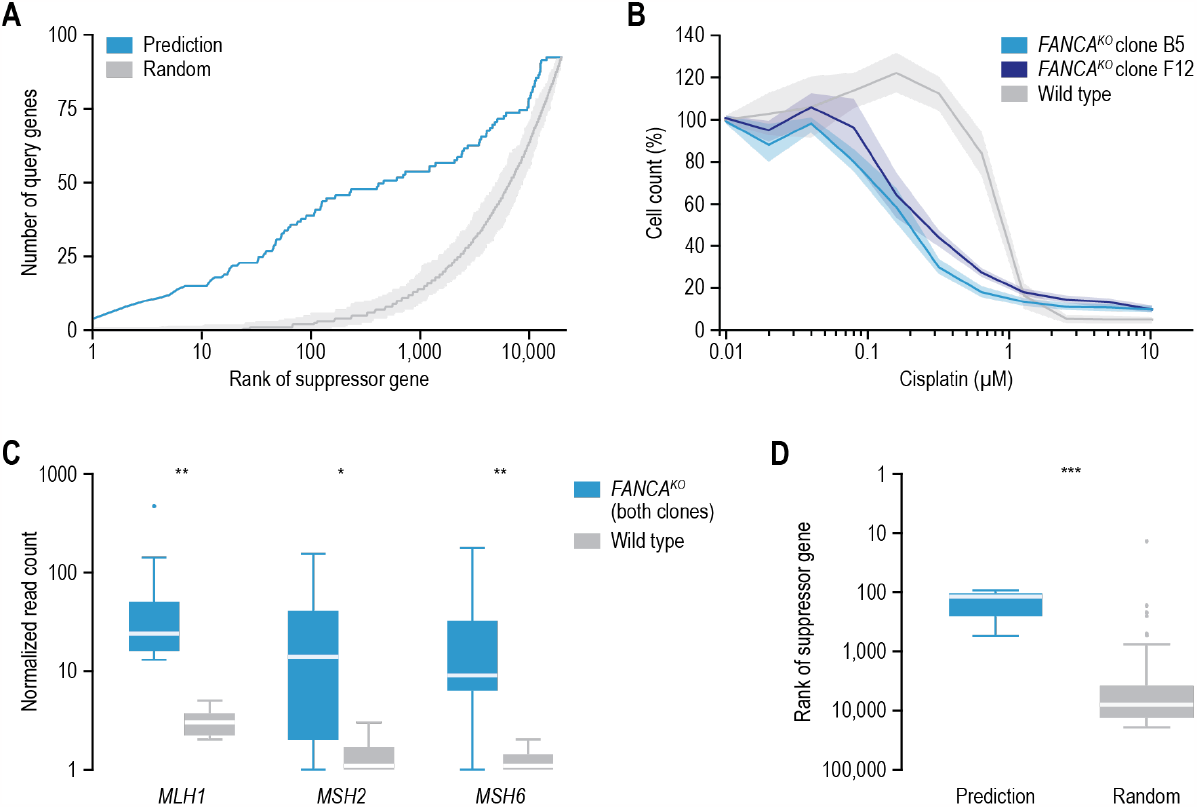
Suppressor gene prediction and validation. (**A**) A suppressor gene prediction model was developed using a random forest classifier. For each query gene, the rank of the validated suppressor gene(s) was determined on both a random gene list and on a list of genes ranked by the likeliness of being a suppressor gene using the prediction algorithm. The rank of the validated suppressor gene was plotted against the number of query genes that interacted with a suppressor gene with that rank. (**B**) *FANCA* knockout cells are sensitive to cisplatin. The indicated cell lines were treated with cisplatin for six days, after which cells were counted. Shading represents the standard error of the mean of at least three independent biological replicates. (**C**) Experimentally identified suppressor genes of *FANCA*. Boxplot showing the normalized read count for guide RNAs targeting the indicated suppressor genes after 18 days of incubation with a concentration of cisplatin that inhibits proliferation by ∼80%. Knockout of the indicated genes specifically suppresses the proliferation defect of *FANCA* knockout cells. (**D**) Comparison of the median rank of confirmed suppressor genes, either in a list of genes ranked by the likeliness of being a suppressor gene using the random forest classifier or in a random gene list. Statistical significance was determined using Mann-Whitney U tests. * p<0.05, ** p<0.005, *** p<0.0005. Horizontal lines in boxplots: median.

Because our literature curated suppression network is not saturated, additional suppressor genes may exist for the 93 query genes in our dataset, that may have been correctly predicted by our random forest model, but not described in the literature. To test this possibility, we experimentally isolated suppressors of *FANCA* in human cells. Loss-of-function mutations in *FANCA* cause Fanconi anemia, a genetic disorder characterized by bone marrow failure and a predisposition to cancer [40]. None of the top 100 suppressor genes that we predicted for *FANCA* had been described in the biomedical literature (**Data S2, S4**). To map *FANCA* suppressors experimentally, we first used CRISPR-Cas9 to create two stable knockout cell lines that both carried a different frameshift deletion in *FANCA* (see Methods). Cells lacking *FANCA* proliferate normally under standard cell culture conditions, but as FANCA is involved in interstrand crosslink repair [41], *FANCA* knockout cells are sensitive to the crosslinking compound cisplatin (**Fig. 7B**). We used a genome-wide CRISPR-Cas9 guide RNA library to identify knockout mutants that could rescue the proliferation defect of the *FANCA* knockout cell lines in the presence of cisplatin. In total, three out of 18,036 tested genes, *MLH1, MSH2*, and *MSH6*, substantially improved proliferation in the presence of cisplatin of both *FANCA* knockout mutants, but not of wild-type cells (**Fig. 7C, Data S5**). The ranks of these three validated suppressor genes on a list of genes ranked by the likeliness of being a suppressor gene using the random forest classifier are significantly lower than what would be expected by chance, with one gene (*MLH1*) ranking in the top 100 (**Fig. 7D, Data S4**, p<0.005, Mann–Whitney U-test). *MLH1, MSH2*, and *MSH6* all encode mismatch repair proteins that can recognize interstrand crosslinks but cannot repair them [42]. In the absence of *FANCA*, expression of these genes thus possibly causes futile DNA repair cycles that may prevent interstrand crosslink repair by other pathways or trigger apoptosis. These experimentally validated suppressor genes show the quality of our predictions, and suggest that our random forest model performs better than can be estimated based on the current set of literature curated suppressors alone. We expect that this predictor will empower future focused searches for suppressor genes in patient populations or cellular models of disease by reducing the number of potential suppressor candidates to a testable number.

## DISCUSSION

We collected 932 suppression interactions from the biomedical literature and used this dataset to define general properties and mechanistic classes of suppression. We found that suppression interactions often linked functionally related genes. General compensation mechanisms were also frequent and tended to affect gene expression or stress response signaling. Furthermore, using *CFTR* and *HBB* as examples, we showed that systematic analysis of suppression interactions can highlight differences in disease pathology.

We discovered that in the absence of a query mutation, suppressor variants are likely deleterious (**Fig. 2**). This suggests that at least some suppressor variants are presumably rare in natural populations and will therefore be difficult to detect using association studies. Nonetheless, suppressor variants may exist for the associated disease alleles and may be identified using alternative methods, such as *in vitro* studies. We have shown here that suppression interactions observed in patients had similar general properties as those found in cultured human cells. For example, both datasets displayed similar fractions of functionally related gene pairs or general suppressor mechanisms (**Fig. 4B, S3B**,**C**). Furthermore, we found a significant overlap between interactions detected in patients and those identified in cultured cells. Together, these observations suggest that cultured cells can be used to discover clinically relevant suppressor genes.

Although many of the general properties we identified for human suppression interactions overlapped with those we previously observed in the budding yeast [8], there were several differences between the two species. Interactions discovered in yeast occurred more frequently between functionally related genes compared to human and were depleted for general compensatory mechanisms, especially those involving signaling or stress response processes such as apoptosis or the immune response (**Fig. 4A**). Suppression in human cells or patients thus often involved more indirect mechanisms of suppression that were not available in unicellular organisms such as yeast. As both the yeast and human datasets are based on literature curated data that can come from specific hypothesis-driven experiments, the datasets may be biased. This bias could differ between the two datasets, due to diverse interests of communities studying different organisms. Nonetheless, we previously mapped a systematic, unbiased experimental suppression network in yeast and showed that the properties of the unbiased network were largely comparable to the literature curated network [8]. Using for example CRISPR-Cas9 knockout screens, a similar unbiased experimental suppression network could be mapped for human genes. Such a network could serve to validate the prevalence of the mechanisms of suppression identified here. Furthermore, as 80% of the described interactions were unique to the suppression network, expanding the human suppression network will reveal new functional connections between genes.

We used the various properties of suppression interactions to develop predictive models of suppression (**Fig. 7, S5**). Although these models can be used to predict suppressor genes for any query gene, the quality of the predictions will be dependent on the availability of functional data for the query and suppressor genes. While the random forest model could also predict more distal functional and general suppression relationships, the majority (∼70%) of interactions in which the validated suppressor ranked in the top 100 of those predicted still involved query-suppressor gene pairs that encode members of the same protein complex or pathway, whereas only ∼15% of all suppression interactions in our dataset occurred between complex or pathway members. The availability of larger, unbiased suppression interaction networks between human genes would likely further improve the prediction of suppression interactions beyond same complex or pathway relationships.

Our dataset lists protective modifiers for most common “monogenic” genetic diseases, including sickle cell disease, β-thalassemia, cystic fibrosis, Huntington’s disease, Duchenne’s muscular dystrophy, and spinal muscular atrophy. For hemoglobinopathies and spinal muscular atrophy, the protective modifiers can completely reverse disease symptoms and have led to the development of effective therapies targeting the suppressor gene [4, 43-45]. Given that suppressor variants have been detected for most common Mendelian diseases and that suppressors can be isolated for >95% of deleterious point mutations in model organisms (our unpublished results), compensatory mutations may exist for nearly all disease alleles. Identification of such suppressor variants may reveal the molecular mechanisms underlying the disease and has the potential to pinpoint new avenues of therapeutic intervention.

## METHODS

### Literature curation

Papers describing potential suppression interactions were collected from multiple sources. First, the *Homo sapiens* “synthetic rescue” and “dosage rescue” datasets were downloaded from the BioGRID on April 11th, 2020 (version 3.5.184) [16]. After removing interactions that did not occur between two human genes, this dataset consisted of 36 genetic interactions described in 21 publications. Second, on April 29th, 2020, the OMIM dataset [17] was downloaded and filtered for entries containing the word “modifier”, which led to the identification of 36 papers potentially describing suppression interactions. Third, we performed PubMed searches for the terms “positive modifier”, “protective modifier”, “synthetic rescue”, “dosage rescue”, “genetic suppression”, and “modifier locus”. Finally, we included papers containing potential suppression interactions that were cited within the examined papers. In total, this resulted in a set of 2,400 papers for further curation (**Data S1**).

All 2,400 papers were read in detail by at least two people. We collected suppression interactions from two types of studies: (i) interactions identified through genetic modifications in cultured human cells and (ii) interactions found through association studies in patients with diseases other than cancer. Two genes were considered to have a suppression interaction when genetic perturbation of a “query” gene led to a disease, reduced survival, decreased cellular proliferation, or was otherwise associated with decreased (cellular) health, which was at least partially rescued by mutation of a “suppressor” gene.

In total, 469 papers were found to describe suppression interactions. From each interaction, we annotated the type of study in which the interaction was identified (cell culture or patients), the query and suppressor genes and mutations and whether these had a loss- or gain-of-function effect, the used cell line or affected tissue, the relative effect size of the suppression, whether any drugs were used, and the disease. All gene names were updated according to the latest approved human gene nomenclature rules [46]. For GWAS, we generally assigned the gene that was closest to the most significant protective SNP as the suppressor gene. However, when data was provided within the paper supporting that another gene was causal, we based our suppressor annotation on this additional evidence. In the case of suppression of *HBB* by SNPs in the intergenic *HBS1L*-*MYB* locus, we assigned all significant SNPs within this locus to *MYB*, which was identified as the causal suppressor gene [47, 48]. Furthermore, all deletions in the HBA locus, for which it often was not specified whether *HBA1* and/or *HBA2* were deleted, were assigned to *HBA2*.

Interactions identified in high-throughput screens yielding >50 suppression interactions were excluded (PubMed IDs 28319085, 32694731, 29891926, and 34764293), as due to their size, these studies would have a disproportionate influence on the complete dataset. For paper 29891926, three interactions that were validated individually were included in the dataset. Also suppression interactions that were intragenic, occurred between more than two genes, or involved the major allele of either the query or the suppressor gene were excluded from the final dataset. In total, the resulting network encompassed 484 different genes and 932 suppression interactions, of which 476 were unique interactions (**Data S2**). Unless indicated otherwise, suppression interactions for *CFTR* and *HBB* were excluded from subsequent analyses, to prevent potential bias introduced by the high number of interactions described for these two query genes.

### Loss-of-function tolerance

The loss-of-function tolerance of query and suppressor genes was evaluated using multiple datasets (**Fig. 2, S1**). First, we used the probability of loss-of-function intolerance of genes that was previously determined based on the frequency of deleterious variants affecting the genes in the human population (gnomad v2.1.1.) [24]. Second, we used the median effect of gene knockout on cell proliferation across a panel of 1,070 cell lines that was determined as the change in abundance of guide RNAs targeting a gene in pooled CRISPR-Cas9 knockout screens [25] (version 22Q1). Third, we used PANTHER version 16.1 to detect the presence of gene orthologs across species [49]. Finally, we considered the number of diseases that were associated with a gene in DisGeNET v7.0 [26]. The phenotypes of query and suppressor genes were compared to those of all other human genes.

### Analysis of gene function and functional relatedness

For analysis of suppression interactions within and across different biological processes (**Fig. 3A, S3A**,**C**), genes were manually assigned to broadly defined functional categories (**Data S2**). Highly pleiotropic or poorly characterized genes were excluded from the analysis. Also interactions involving query genes annotated to the “Protein folding & glycosylation” class were removed from consideration, as only one interaction fell into this category. G:Profiler version e107_eg54_p17_bf42210 [50] was used to identify suppressor genes within the broader “Signaling & stress response” category that had a role in protein phosphorylation (GO:0006468) or apoptosis (GO:0006915).

We used systematic, genome-wide datasets describing protein localization, GO term annotation, co-expression, protein complex membership, and pathway membership to evaluate the functional relatedness between query-suppressor gene pairs (**Fig. 3B-C, S3B**). In each case, only gene pairs for which functional data was available for both the query and the suppressor gene were considered. Protein localization was determined based on immunofluorescence staining data available in The Human Protein Atlas version 21.1 [51]. Two proteins were considered to co-localize if they were found in at least one shared cellular compartment. GO co-annotation was calculated based on biological process terms with less than 500 annotated genes. Co-expression data was derived from SEEK [52] as explained previously [53]. Proteins that were annotated to the same protein complex in either CORUM 4.0 [54] or BioPlex 3.0 [55] were considered as co-complexed. Proteins in distinct non-overlapping protein complexes were considered not co-complexed. The same approach was used to define co-pathway membership using Reactome data (downloaded January 2020) [56]. For each of these datasets, we calculated the overlap with the suppression interactions. The expected overlap by chance was calculated by considering all possible pairs between a background set of queries and suppressors. The background set of queries consisted of genes found as queries in the suppression network. As a background set of suppressors, we considered all genes in the genome. Pairs with a suppression interaction were removed from the background set. For a given functional standard, we defined as fold enrichment the ratio between the overlap of suppression gene pairs and the overlap of the background set with that standard. Significance of the overlap was assessed by Fisher’s exact tests. To evaluate the functional relatedness of multiple suppressors of the same query gene (**Fig. 3C**), only suppressors described in different papers were considered.

### Overlap with other types of interactions

We compared our suppression interaction network to three different interaction datasets collected from the BioGRID [16]: physical interactions, negative genetic interactions, and positive genetic interactions (**Fig. S2**). For the genetic interaction datasets, we removed papers from the BioGRID data that were used for the suppression interaction literature curation. Overlap of the interaction networks with our suppression interaction dataset were calculated as explained above (see “**Analysis of gene function and functional relatedness**”).

To investigate whether suppressor genes with a role in transcription or chromatin organization affected expression of members of the same pathway as the query gene, we used transcription factor (TF) target gene information from g:Profiler e106_eg53_p16_65fcd97 [50] and MotifMap [57] and pathway annotations from Reactome [56]. To exclude non-specific annotations, we only considered pathways and g:Profiler TF-target lists with less than 100 members. MotifMap and g:Profiler gave comparable results, with respectively 38% and 50% of the suppressor genes that encode transcription factors with known targets affecting expression of corresponding query pathway members.

### Mechanistic classes

Suppression interactions were assigned to distinct mechanistic classes (**Fig. 4, 5C**). Gene pairs that had the same biological process annotation (**Data S2**) or gene pairs that were not annotated to the same biological process but encoded members of the same complex or pathway (see “**Analysis of gene function and functional relatedness**” for details on the used datasets) were considered to be functionally related. These functionally related gene pairs were further subdivided into subclasses. First, gene pairs that encoded subunits of the same protein complex were assigned to the “Same complex” subclass. Second, gene pairs that encoded members of the same pathway were assigned to the “Same pathway” subclass. Third, gene pairs that shared a biological process annotation but functioned in different pathways were assigned to the “Alternative pathway” subclass. Finally, all other functionally related gene pairs were assigned to the “Uncharacterized functional relation” subclass.

Gene pairs that did not have a functional relationship were further subdivided based on the function of the suppressor gene. Suppressor genes that were annotated to the biological processes “Transcription & chromatin organization”, “Translation & RNA processing”, “Protein degradation”, and “Signaling & stress response” were assigned to the corresponding subclasses. The remaining gene pairs were assigned to the “Other/unknown” class.

### Co-occurrence of mutations in cancer models and patients

To determine whether the effect of knockout of a given gene on cellular fitness strongly depended on the genetic background, we examined fitness data from genome-scale CRISPR-Cas9 gene knockout screens across 1,070 cancer cell lines from the DepMap project [25]. Because the variance in gene knockout fitness varied depending on the average fitness of the gene knockout across cell lines, we fitted a quadratic model to the fitness data and used it to determine whether a given gene had a higher fitness variance across cell lines than expected (**Fig. S4A**,**B**).

To evaluate the frequency of co-occurrence of mutations in query-suppressor gene pairs (**Fig. 6**), we used data from two sources. First, we used cell line mutation data from the Cancer Cell Line Encyclopedia from DepMap [37]. We considered only “damaging mutations” as defined by DepMap [37]. Second, we examined the co-occurrence of mutations in tumor samples collected from 69,223 patients across a curated set of 213 non-overlapping studies on cBioPortal [38]. We excluded variants of unknown significance as defined by cBioPortal [38]. We then calculated how often the query and corresponding suppressor genes were co-mutated compared to a background set of gene pairs, as explained above (see “**Analysis of gene function and functional relatedness**”), with the exception that for cBioPortal analysis, we used genes found as suppressor genes in the suppression interaction dataset of interest as background set.

### Predicting suppressor genes

We predicted potential suppressor genes by ranking all genes in the human genome by their functional relationship to the query gene (**Fig. 7A, S5, Data S3, S4**). We used two different models to do this. First, based on a suppressor-prediction algorithm we previously developed for yeast [5], we evaluated the following functional relationships in this order of priority: co-complex (highest priority), co-pathway, co-expression, and co-localization (lowest priority). Thus, genes with co-complex relationships were ranked above those with only co-pathway relationships. Additionally, the order between genes within a given set was established by evaluating the rest of the functional relationships. For instance, the set of genes that were co-expressed with the query gene, but did not encode members of the same complex or pathway, was further ranked by whether the encoded protein co-localized (highest rank) or not (lowest rank) with the query protein. Second, we used the same four functional datasets, genetic interactions, protein-protein interactions, and co-mutation data in cancer cell lines to train a random forest classifier using the R package “randomForest” [58]. The complete set of suppression interactions, including those described for *CFTR* and *HBB*, was used to train the model. Performance of the predictor was evaluated with out-of-bag samples. See the previous sections for details on the used datasets.

### Experimental validation of predicted suppressor genes

HEK293T cells were maintained in Dulbecco’s modified Eagle’s medium (DMEM; ThermoFisher) supplemented with 10% FBS, 100 U/mL penicillin, and 100 mg/mL streptomycin. HAP1 cells were cultured in Iscove’s Modified Dulbecco’s Medium with GlutaMAX (IMDM; ThermoFisher) supplemented with 10% FBS, 100 U/mL penicillin, and 100 mg/mL streptomycin. All cells were cultured in humidified incubators at 37° C and 5% CO2.

TKOv3 library lentivirus was produced by cotransfection of lentiviral vectors psPAX2 (packaging vector; Addgene #12260) and pMDG.2 (envelope vector; Addgene #12259) with the TKOv3 lentiCRISPR plasmid library [59]. Briefly, HEK293T cells were seeded at a density of ∼7 × 10^6^ cells per 15-cm dish and incubated overnight, after which cells were transfected with a mixture of psPAX2 (4.8 μg), pMDG.2 (3.2 μg), TKOv3 plasmid library (8 μg), and X-tremeGene 9 (48 μL; Roche), in accordance with the manufacturer’s protocol. At 24 hours after transfection, the medium was changed to serum-free, high BSA growth medium (DMEM, 1% bovin serum albumin, 100 U/mL penicillin, and 100 mg/mL streptomycin). Virus-containing medium was harvested 48 hours after transfection, centrifuged at 1500 rpm for 5 min, and stored at -80° C. Functional titers in HAP1 cells were determined by infecting cells with a titration of TKOv3 lentiviral library in the presence of polybrene (8 μg/mL). At 24 hours after infection, medium was replaced with puromycin (1 μg/mL) containing medium to select for transduced cells, and incubated for 48 hours. The multiplicity of infection (MOI) of the titrated virus was determined 72 hours after infection by comparing the percent survival of infected cells to non-infected control cells.

To create stable cell lines lacking *FANCA*, guide RNA (gRNA) 5’-TACCACATCCACTCACCCTG-3’ targeting *FANCA* was cloned into pLentiCRISPRv2-Blast-mU6. This vector is based on lentiCRISPRv2-Blast (Addgene #83480), which carries both the Cas9 enzyme and a gRNA expression cassette, in which the hU6 promoter driving the gRNA cassette has been replaced with the mU6 promoter from plasmid pmU6-gRNA (Addgene #53187). Lentivirus was produced using the procedure described above, and 24 hours after transduction, transduced HAP1 cells were selected with 25 μg/mL blasticidin for 72 hours, followed by single clone isolation. Gene editing was confirmed by PCR amplification of the targeted region using primers 5’-TACACTCTCTCGTCGCCGCACA-3’ and 5’-CAGGTTCCGGGCAGGTAGGGAA-3’, followed by Sanger sequencing and TIDE analysis [60]. *FANCA*^*KO*^ clone B5 carries a 7 basepair deletion at residue 361 (361del7) and *FANCA*^*KO*^ clone F12 carries a single basepair deletion at the same location (361delG), based on CDS NM_000135.4. Both *FANCA*^*KO*^ cell lines spontaneously diploidized during the single cell isolation, and are homozygous for the edits in *FANCA*. Because the *FANCA* knockout clones were diploid, diploidized HAP1 cells were used as wild-type controls in all further experiments.

For proliferation assays with cisplatin (**Fig. 7B**), 1,000 cells were seeded in 96-wells plate. The following day, cisplatin was added at 2-fold serial dilutions and medium containing cisplatin was renewed after three days. Six days after the initiation of treatment, cell counts were measured using a Cytation 5 Cell Imaging Multimode Reader (Agilent BioTek).

To identify suppressor genes that could rescue the proliferation defect of *FANCA* knockout cells in the presence of cisplatin (**Fig. 7C**), a total of 50 × 10^6^ cells per cell line were infected with the TKOv3 lentiviral library (71,090 gRNAs) at an MOI of ∼0.3 to achieve ∼200-fold coverage of the library after selection. At 24 hours after infection, medium was replaced with puromycin (1 μg/mL) containing medium to select for transduced cells. Two days later, selected cells were split into two technical replicates containing 15 × 10^6^ cells each, treated with cisplatin, and passaged every 3 days. Wild-type diploidized HAP1 cells were treated with 1.25 μM cisplatin and the *FANCA* knockout cell lines, that show increased sensitivity to crosslinking agents, were treated with 0.3 μM cisplatin. A total of 20 × 10^6^ cells were collected for genomic DNA extraction at 0 and 18 days after puromycin selection. Genomic DNA was extracted from cell pellets using the QIAamp Blood Maxi Kit (Qiagen), according to the manufacturer’s instructions. Guide RNA inserts were amplified via PCR using primers harboring Illumina TruSeq adapters i5 and i7 barcodes at 50-fold coverage, and the resulting libraries were sequenced on an Illumina HiSeq4000 instrument.

Sequencing reads were assigned to gRNAs using the MAGeCK count module [61]. Read counts were normalized to the total number of reads per sample (**Data S5**). Note that a high variability in read counts between gRNAs targeting the same gene is expected under conditions of extreme positive selection. Guide RNAs that were absent in 2 or more of the day 0 samples were removed from consideration. Genes were called suppressors if the normalized read count across clones, technical replicates, and gRNAs targeting the gene was significantly higher at the end of the screen (day 18) in *FANCA* knockout cells compared to wild-type cells (p<0.05 Wilcoxon test). Furthermore, to exclude genes that only had a minor effect on proliferation, we required that at least 50% of the gRNAs and replicates per gene had a log_2_ fold change>2 between day 18 and day 0 samples in both *FANCA* knockout clones but not in the wild type control. These thresholds led to the identification of three high-confident suppressor genes: *MLH1, MSH2*, and *MSH6*.

## SUPPLEMENTARY DATA FILES

**Data S1. List of papers that were analyzed**.

This file lists the PubMed IDs of all the papers that were read during the literature curation.

**Data S2. Suppression interactions curated from the literature**

This file contains the suppression interactions that met our selection criteria (see Methods), along with the corresponding PubMed IDs, details on the query and suppressor genes and mutations, and information on the conditions under which the interactions were identified.

**Data S3. Suppressor gene predictions using the functional prioritization model**

For each of the 93 query genes in our dataset, all genes in the human genome were ranked based on their likeliness of being a suppressor gene using a functional prioritization model. Listed are query gene name, suppressor gene name, prediction score, and prediction rank. Prediction scores range from 0 to 1111. Higher prediction scores or lower prediction ranks correspond to a higher likeliness of a gene to be a suppressor of the given query gene.

**Data S4. Suppressor gene predictions using the random forest classifier**

For each of the 93 query genes in our dataset, all genes in the human genome were ranked based on their likeliness of being a suppressor gene using a random forest classifier. Listed are query gene name, suppressor gene name, prediction score, and prediction rank. Prediction scores range from 0 to 1. Higher prediction scores or lower prediction ranks correspond to a higher likeliness of a gene to be a suppressor of the given query gene.

**Data S5. Experimental validation of predicted suppressor genes**

Experimental identification of suppressor genes of *FANCA* using CRISPR-Cas9 knockout screens. This file contains normalized read counts for 71,090 gRNAs across 12 samples: two timepoints (0 and 18 days of cisplatin treatment) of one wild-type and two *FANCA* knockout cell lines each screened in duplicate.

## DECLARATIONS

### Ethics approval and consent to participate

Not applicable

### Consent for publication

Not applicable

### Availability of data and materials

All data generated or analyzed during this study are included within the article and its supplementary files.

### Competing interests

The authors declare that they have no competing interests.

### Funding

This work was supported by an Eccellenza grant from the Swiss National Science Foundation (PCEGP3_181242) (J.v.L) and a Ramon y Cajal fellowship (RYC-2017-22959) (C.P.).

## Acknowledgements

We thank Claire Paltenghi for critical reading of the manuscript.

## Notes

### Competing Interest Statement

The authors have declared no competing interest.

### Summary of Updates

In this version of the manuscript, we have added experimental validations of our suppressor gene predictions.

