## Supplementary Figures for "Global analysis of suppressor mutations that rescue human genetic defects"

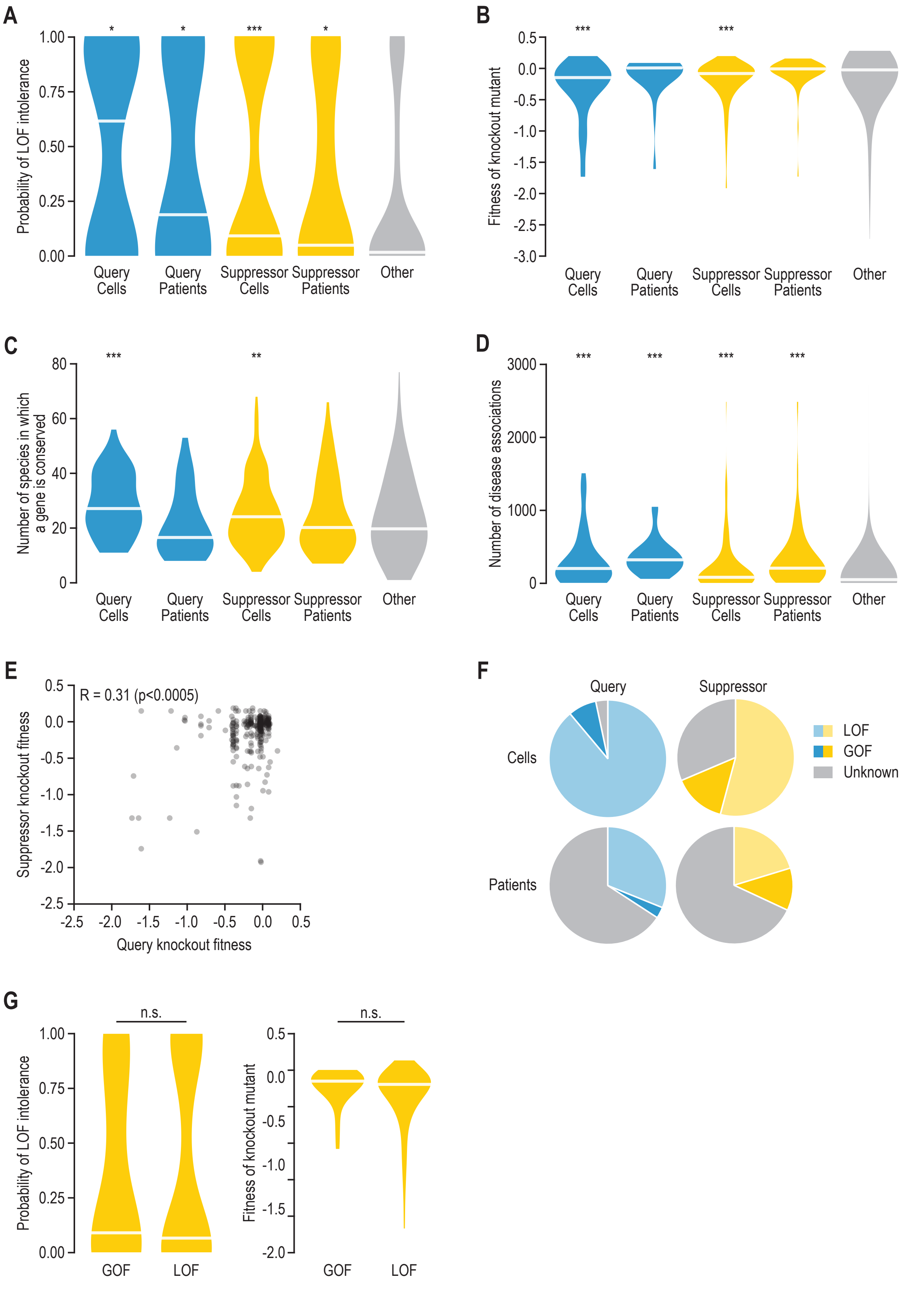


**Fig. S1. Suppressor genes are important for maintaining health and cellular fitness.** (**A**) Probability of loss-of-function intolerance for query genes, suppressor genes, and all other genes, based on the frequency of deleterious variants affecting the genes in the human population [24]. Query and suppressor genes were further subdivided based on whether the interactions had been described in cultured cells or patients. (**B**) Median effect of gene knockout on cell proliferation determined as the change in abundance of guide RNAs targeting a gene in pooled CRISPR-Cas9 screens across 1,070 cell lines [25], for the same gene groups as in (A). (**C**) Number of species in which an ortholog of the query or suppressor gene is present. (**D**) The number of diseases that are associated with a gene in DisGeNET [26], for the same gene groups as in (A). (**E**) Median effect of gene knockout on cell proliferation as in (B) for query-suppressor gene pairs. The Pearson correlation coefficient and corresponding p-value are indicated. (**F**) The fraction of query and suppressor genes that have loss-of-function (LOF), gain-of-function (GOF), or unknown modes of action. (**G**) Probability of loss-of-function intolerance and the effect of gene knockout on cell proliferation as in (A) and (B), for suppressor genes that carry loss-of-function or gain-of-function mutations. n.s. = not significant. Statistical significance compared to the “Other” group (A-D) or between LOF and GOF groups (G) was determined using Mann-Whitney U tests. * $p<0.05$, ** $p<0.005$, *** $p<0.0005$. Horizontal lines in violin plots: median.

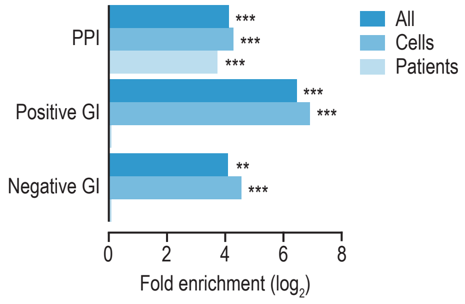


**Fig. S2. Overlap with other interaction networks.** Fold enrichment for overlap of suppression interactions with protein-protein interactions (PPI), or with positive and negative genetic interactions (GI), either for all suppression interactions, or for interactions identified in cultured cells or in patients only. Fisher's exact tests were performed to determine statistical significance of the results. * $p<0.05$, ** $p<0.005$, *** $p<0.0005$.

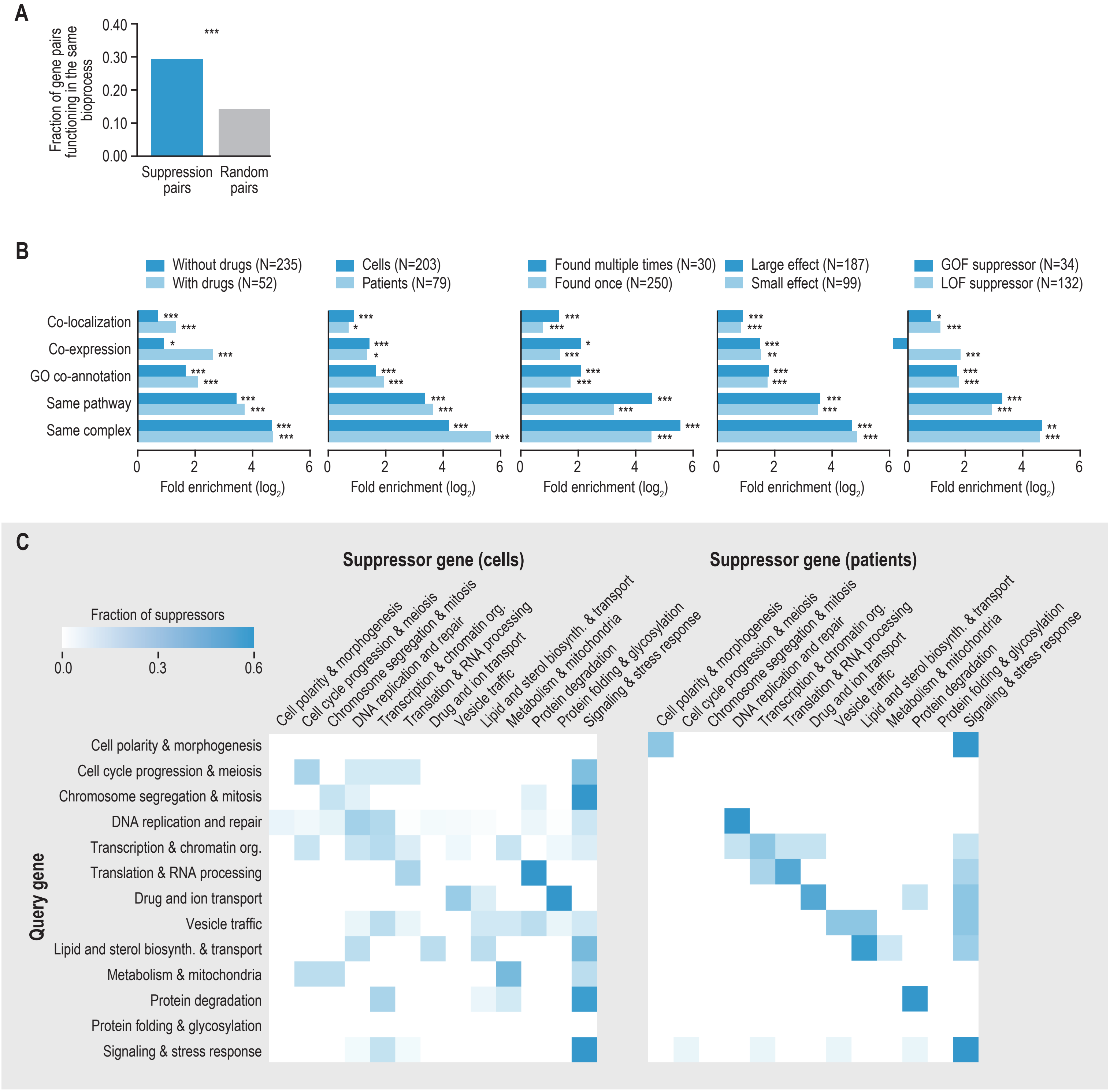


**Fig. S3. Functional connections between query and suppressor genes.** (**A**) Fraction of query-suppressor or random gene pairs sharing a biological process annotation. (**B**) Fold enrichment for co-localization, co-expression, GO co-annotation, same pathway membership, and same complex membership for different subsets of suppression interactions. The number of interactions in each subset is indicated in the legends. Note that the total number of interactions varies per analysis, because some interactions may have neither or both annotations (i.e. an interaction may have been identified both in the presence and in the absence of a drug). (**C**) Frequency of suppression interactions connecting genes within and across indicated biological processes for interactions identified in cultured cells (left) or in patients (right). Color reflects the fraction of suppressor genes belonging to a particular biological process for all interactions involving query genes annotated to a given biological process. Fisher's exact tests were performed to determine statistical significance of the results in (A) and (B). * $p<0.05$, ** $p<0.005$, *** $p<0.0005$.

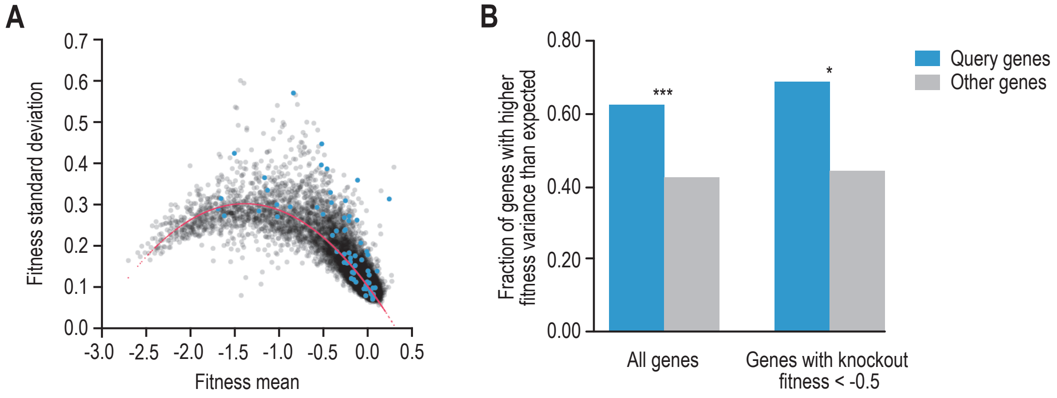


**Fig. S4. Query gene knockout is associated with large variation in fitness across cell lines.** (**A**) Plotted are the mean fitness of a knockout mutant across 1,070 cell lines against the fitness standard deviation [25]. Each data point represents a gene, query genes are highlighted in cyan. Pink = the model that was fit to the data. (**B**) Fraction of genes that have a higher variance in fitness across cell lines than expected by chance given the model shown in (A), for either all genes or for genes with an average knockout fitness across cell lines <-0.50. Statistical significance was determined using one-sided Fisher's exact tests. * p<0.05, ** p<0.005, *** p<0.0005.

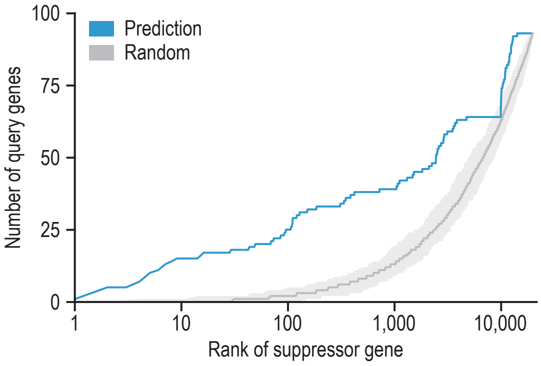


**Fig. S5. Suppressor gene prediction.** A suppressor gene prediction model was developed based on the strong functional connection between query and suppressor genes, similar to a model we previously developed for yeast [5]. For each query gene, the rank of the validated suppressor gene(s) was determined on both a random gene list and on a list of genes ranked by the likeliness of being a suppressor gene using the prediction algorithm. The rank of the validated suppressor gene was plotted against the number of query genes that interacted with a suppressor gene with that rank.
